# Allelic Diversity and Core Conservation of Type III Effectors Across Xanthomonads Causing Bacterial Spot of Pepper and Tomato

**DOI:** 10.64898/2026.08.04.742720

**Authors:** Apekshya Parajuli, Aastha Subedi, Amandeep Kaur, Sophia McDuffee, Gerald V. Minsavage, Fernanda Iruegas Bocardo, Jeannie M. Klein-Gordon, Anuj Sharma, Gary E. Vallad, Erica M. Goss, Jeffrey B. Jones

## Abstract

Bacterial spot of tomato and pepper (BST/P) is an economically devastating disease caused by four distinct *Xanthomonas* pathogens: *X. euvesicatoria* pv. *euvesicatoria* (*Xe*), *X. euvesicatoria* pv. *perforans* (*Xp*), *X. hortorum* pv. *gardneri* (*Xg*), and *X. vesicatoria* (*Xv*). A key component of virulence in these pathogens is the type III secretion system (T3SS), which delivers type III effector (T3E) proteins into host plant cells. To comprehensively characterize T3E repertoires and assess the stability of core effectors at a population scale, we evaluated a global dataset comprising 1,037 quality-filtered genomes, including 585 *Xp*, 350 *Xe*, 69 *Xg*, and 33 *Xv* strains. Across this collection, genes for six effectors were present in 100% of the examined genomes (XopK, XopL, XopM, XopN, XopX, and XopZ1) and an additional four effectors in ≥95% of genomes (XopK, XopL, XopM, XopN, XopX, and XopZ1). *Xp* and *Xe* populations maintained large total effector repertoires with extensive allelic variation, displaying exceptional polymorphism within XopD and XopAD. In contrast, *Xg* and *Xv* exhibited highly stable effector profiles with markedly reduced allelic diversification across geographic regions and decades. Disruptive mutations, including early stop codons and frameshifts mutations, in genes for XopAZ, XopAF, and XopAR were prevalent across specific pathogens pointing to ongoing pseudogenization and targeted gene loss. These findings provide a high-resolution characterization of the conserved and variable components of the BST/P pathogen effector arsenal and serve as a foundation for monitoring population evolution and breeding durable disease resistance to multiple pathogens.

## Introduction

Prokaryotes utilize specialized protein secretion mechanisms to transport proteins from the cytoplasm to host cells (Green & Mecsas, 2016). A critical example is the type III secretion system (T3SS), a macromolecular "injectisome" that delivers bacterial type III secretion effector proteins directly into the host cytoplasm (Rosqvist et al., 1994). The T3SS has a virulence function in pathogens such as *Xanthomonas* and *Pseudomonas* (White et al., 2009; Su et al., 2025), but also facilitates beneficial interactions in symbionts like *Rhizobium* (Zboralski et al., 2022). Type III effectors (T3Es) act as molecular tools to alter the host cell machinery for the pathogen’s benefit. A T3E may contribute to virulence (Kim et al. 2011; Jacobs et al., 2013; Nomura et al., 2006), symptom development (Badel et al., 2006; Kim et al., 2009), pathogenicity (Boureau et al. 2006; Gaudriault et al., 1997), host specificity (Rybak et al., 2009), and ultimately pathogen fitness (Sharma et al. 2021). T3Es can have a wide array of activities, including interfere with the host cell cytoskeleton to promote attachment and invasion, modulate cellular trafficking processes, cytotoxicity, and, critically subvert the host immune system (Deng et al., 2014)

The co-evolutionary interaction between pathogens and their hosts is a dynamic arms race, with the T3SS and its effectors playing central roles. While the T3SS apparatus itself is highly conserved across different pathogens, the number, sequence, and function of the T3Es are highly variable (Chan et al., 2025). T3E repertoires can be conceptualized as having a core set of effectors essential for basal virulence and accessory effectors that may provide functional redundancy or are conditionally required to promote fitness on a specific host or under particular circumstances (Lindeberg et al., 2012). Bacteria can acquire T3Es through horizontal gene transfer mediated by mobile genetic elements like plasmids and pathogenicity islands (Stavrinides & Guttman, 2004) or phages (Hulin et al., 2023). Host organisms have evolved intracellular resistance (R) proteins to recognize specific T3Es, triggering a defense response known as effector-triggered immunity (ETI) (Chan et al., 2025; Jones & Dangl, 2006). Pathogens are under selection to evade this recognition while maintaining virulence, which drives allelic variation (Chan et al., 2025; Dillon et al., 2019; Pritchard & Birch, 2014)

In *Xanthomonas*, T3Es are primarily known as XOPs (*Xanthomonas* outer protein) with a few historical exceptions like AvrBs1, AvrBs2, and AvrBs3. There are sixty-five known XOP families (Chan et al. 2025). This study examines the repertoires and allelic variation of *Xanthomonas* T3Es within the four *Xanthomonas* pathogens responsible for bacterial spot of tomato and pepper (BST/P): *X. euvesicatoria* pv. *euvesicatoria* (*Xe*), *X. euvesicatoria* pv. *perforans* (*Xp*), *X. vesicatoria* (*Xv*), and *X. hortorum* pv. *gardneri* (*Xg*). BST/P have been economically important diseases worldwide for decades (Kyeon et al. 2016, Burlakoti et al. 2018, Klein-Gordon et al. 2021, Subedi et al. 2023a, Parajuli et al. 2024, Chen et al. 2024, Liao et al. 2024, Preangtong et al. 2025, Lue et al. 2010, Rotondo et al. 2022, Schwartz et al. 2015) causing 50% or more yield loss in favorable climates (Bashan et al., 1985; Pohronezny & Volin, 1983; Ritchie, 2000)*Xe* is the major BSP causing pathogen whereas *Xp* is the major BST causing pathogen. In the past, *Xe* was predominantly isolated from diseased tomato plants but in last 4 decades, *Xp* has emerged as a major BST pathogen (Subedi et al. 2023b, Chen et al. 2024, Klein-Gordon et al. 2021), which is partly attributed to presence of bacteriocins in *Xp* that are effective against *Xe* (Jones et al. 1998, Klein-Gordon et al. 2023). *Xv Xg* appears to be more common in cooler tomato production regions (Ma et al. 2011, Khanal et al. 2020, Dia et al. 2022). In recent years *Xv* has been rarely isolated from diseased tomato and pepper (Burlakoti et al. 2018; Chen et al. 2024; Egel et al. 2018). Evidence of selection to avoid R-gene detection includes single nucleotide substitutions in *avrBs2* in *X. euvesicatoria* (Gassman et al, 2000) and multiple different loss of function mutations affecting *avrXv3* in *X. perforans* (Timilsina et al., 2016). Understanding the distribution of T3Es and their allelic variation in *Xanthomonas* populations is crucial for developing durable disease resistance strategies.

Given the economic significance of pathogen strains that overcome host resistance genes, the primary objective of this research is to comprehensively characterize the T3E repertoires of these four pathogens. While Potnis et al. (2011) previously established the baseline for conserved and taxon-specific effectors using one representative of each taxon, this study seeks to determine how these repertoires shift with bigger sample sizes. This research also goes beyond previous work in examining T3E repertoires (Schwartz et al. 2015, Subedi et al 2023b, Chen et al. 2024, Jibrin et al. 2024) by analyzing allelic variation across BST/P pathogen strains isolated around the world. Specifically, we aim to re-evaluate the core effectors to assess their stability, while simultaneously investigating recent evolutionary changes that may have resulted from ongoing adaptation on tomato or pepper hosts. Our results provide new insights into the dynamics of XOPs in the BST/P pathogens, an essential reference dataset for deciphering the molecular basis of virulence, and conserved components of the pathogens’ T3E virulence arsenals for potential development of durable resistance strategies.

## Materials and Methodology

### Genomes used in this study

Genomes representing a diverse collection of four bacterial spot-causing *Xanthomonas* taxa on tomato and pepper: *Xp*, *Xe*, *Xg*, and *Xv* were retrieved from the NCBI database (https://www.ncbi.nlm.nih.gov/). An initial quality assessment was performed on all genome assemblies and only those with less than 500 contigs, sequencing coverage ≥30x, genome completeness above 97% and contamination below 3% were retained for downstream analyses. Average nucleotide identity (ANI) analysis was then performed on all high-quality genomes against the type strains of their respective species *Xp* (ICMP 16690), *Xe* (LMG 27970), *Xg* (ATCC 19865), *Xv* (ATCC35937) using FastANI(v.1.3) with the ANIb algorithm (Jain et al. 2018). Additionally, we sequenced 77 *Xp* and 23 *Xe* new genomes and we re-sequenced six *Xg* strains with previously reported genomes of low quality. The final dataset includes 585 *Xp*, 350 *Xe*, 69 *Xg* and 33 *Xv* genomes (Supplementary Table 1a-d).

### Whole genome sequencing and assembly

For genome sequencing, DNA was extracted from the overnight cultures using the Wizard Genomic DNA Purification Kit (Promega, Madison, WI) following manufacturer’s instructions. DNA samples were sent for paired-end sequencing (2 x 150 bp) on an Illumina NovaSeq platform at SeqCenter (Pittsburgh, PA). Raw reads were quality trimmed, and adapters were removed using Trim Galore (v0.6.10) with default parameters (Krueger, 2015). Filtered reads were then assembled using SPAdes (v3.10.1) with “careful” parameters and *k*-mer lengths of 21, 33, 55, 77, 99, and 127 (Bankevich et al., 2012). Contigs smaller than 500 bp and *k*-mer coverage of 2.0 were filtered out. Final assemblies were polished using Pilon (v1.24) with the default parameters (Walker et al., 2014). All genome statistics were calculated using a python script available at: https://github.com/sujan8765/nepgorkhey_python/blob/master/genome_stats.py. Assembled genomes were submitted to NCBI GenBank database and annotated using the NCBI PGAP pipeline.

### Construction of species-specific T3E reference databases

We constructed T3E reference databases for *Xp*, *Xe*, *Xg* and *Xv* separately. Specifically, the curated T3Es database available at https://euroxanth.ipn.pt/doku.php?id=bacteria:t3e:t3e was first used as a query in tBLASTn searches against all genomes. For *Xp* and *Xe*, effectors were considered present when tBLASTn hits exhibited ≥70% sequence coverage and ≥95% sequence identity. For *Xg* and *Xv,* a more relax*e*d cutoff of ≥50% sequence coverage and ≥50% percentage identity was applied to capture more divergent homologs. All BLAST hits were manually inspected to ensure effector presence, and a single representative sequence per effector was then selected to construct the final taxon-specific reference databases.

### Identification of T3Es and allelic variation

The T3E reference databases were queried against the genomes of each taxon using tBLASTn with an e-value cutoff of 1e-3, ≥95% identity and ≥70% sequencing coverage. Effectors identified were then manually examined to characterize allelic variation. Any sequence variation resulting in an amino acid change in an effector was assigned to a distinct allele type. Effectors that could not be assembled into a single contig, as well as those containing frameshifts or early stop codons, were also assigned separate allele types and categorized as contig breaks (CTB), frameshifts (FS), or early stop codons (ESC). Heatmaps summarizing allelic diversity within each species were generated using Heatmaply (Galili et al., 2018).

### Phylogenetic analysis

For phylogenetic analysis of strains within each species, all genomes were annotated using Prokka (v1.14.6) (Seemann, 2014). The resulting gff files were then used as input for Roary (v1.007002) to generate core gene alignments (Page et al., 2015). These core gene alignments were subsequently used as input for RAxML (v8.2.10) to construct maximum likelihood phylogenies using substitution model GTRGAMMAI with 500 bootstraps (Kozlov et al. 2019). The final phylogenetic trees were visualized using iTOL (v7) (Letunic & Bork, 2021).

## Results

### Geographic Distribution of BST/P Pathogen Genomes

The *Xp* dataset was the most extensive, comprising 586 genomes (1991-2021) predominantly isolated from tomato (n=531) from 15 countries on five continents, though heavily skewed toward the USA (n=474) (Figure 1, Supplementary Table 1a). Two strains (XTN47 and XpT2) originally classified as *Xe* were reclassified as *Xp* based on ANI values (Supplementary Table 2a).. In contrast to *Xp*, the 350 *Xe* genomes (1957–2021) were primarily pepper-associated (Supplementary Table 1b) from 17 countries, including large collections from the USA (n=178) and Taiwan (n=72) (Figure 1, Supplementary Table 1b). The *Xg* collection (1953-2016, n=69) was largely isolated from tomato in the Americas and Africa (Supplementary Table 1c, Figure 1). The *Xv* genomes (1955-2023, n=33) were primarily associated with tomato and had a global distribution (North/South America, Oceania) spanning nearly 70 years (Figure 1, Supplementary Table 1d).

**Figure 1.**
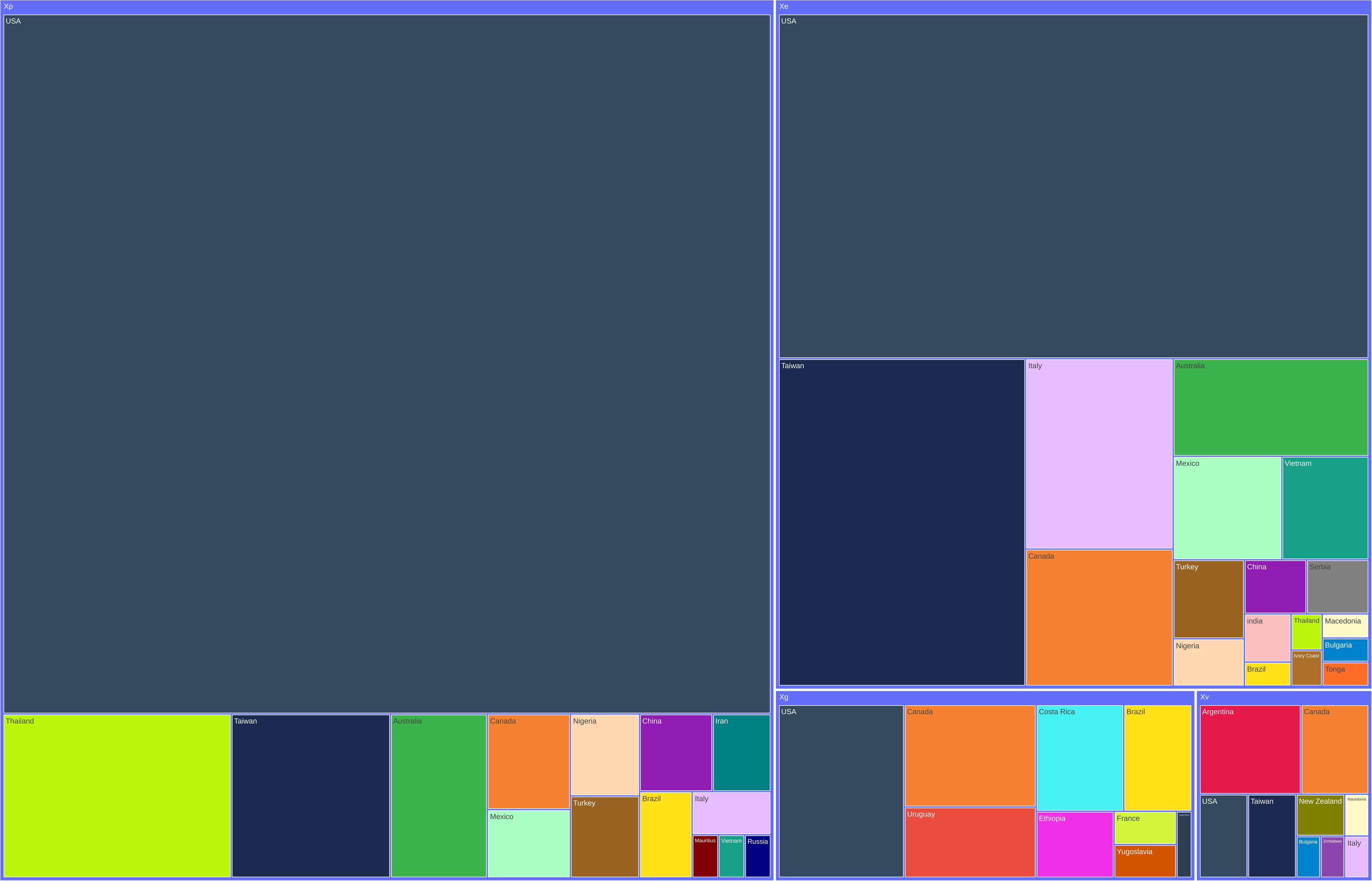
Representation of country of isolation of strains with genomes examined in this study. Xp, Xe, Xg and Xv represents, *Xanthomonas euvesicatoria* pv. *perforans, Xanthomonas euvesicatoria* pv. *euvesicatoria, Xanthomonas hortorum* pv. *gardneri and Xanthomonas vesicatoria* respectively. Size of rectangle is proportional to the number of strains from each location, created with Treemap.

**Figure 2.**
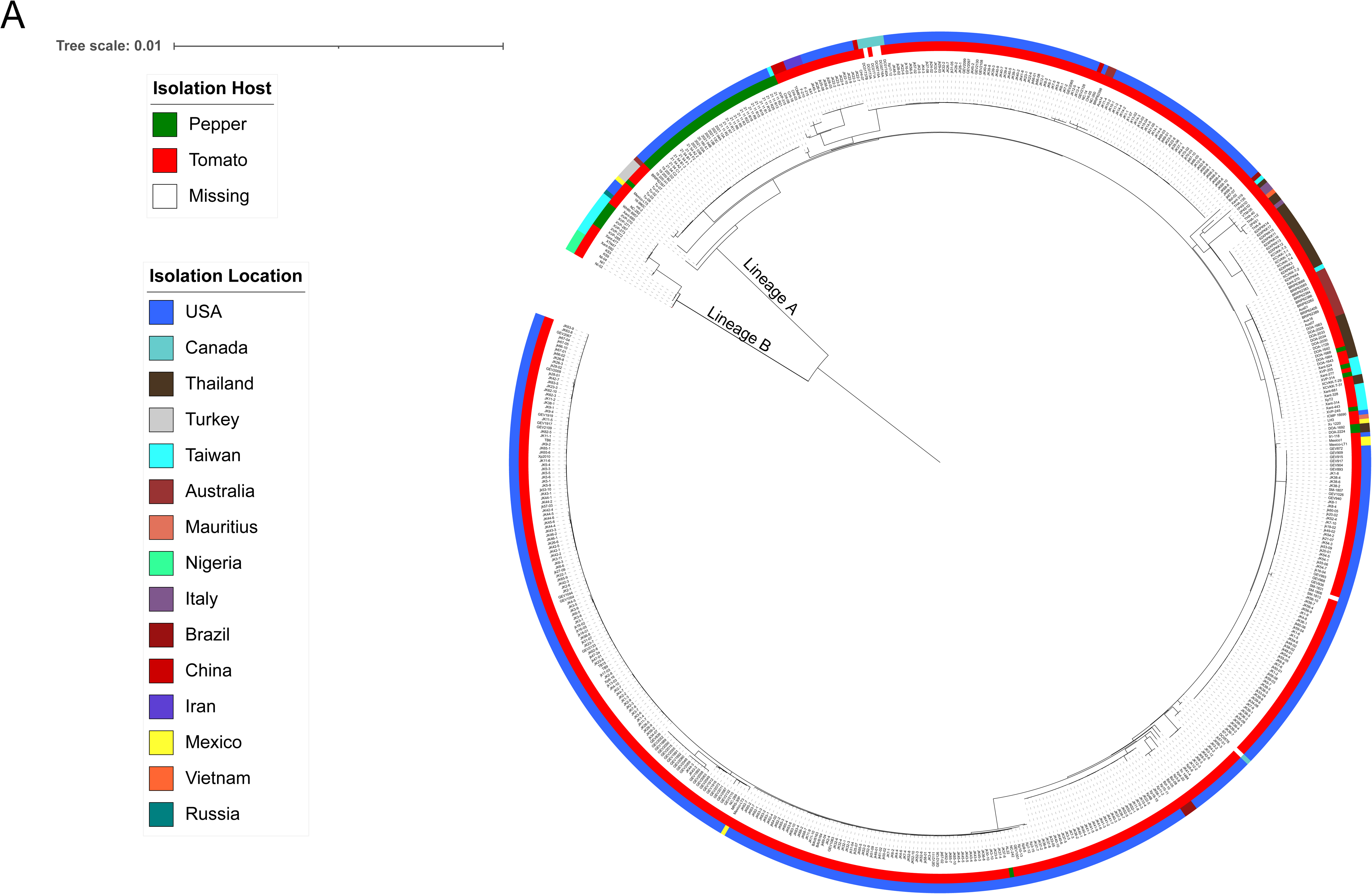
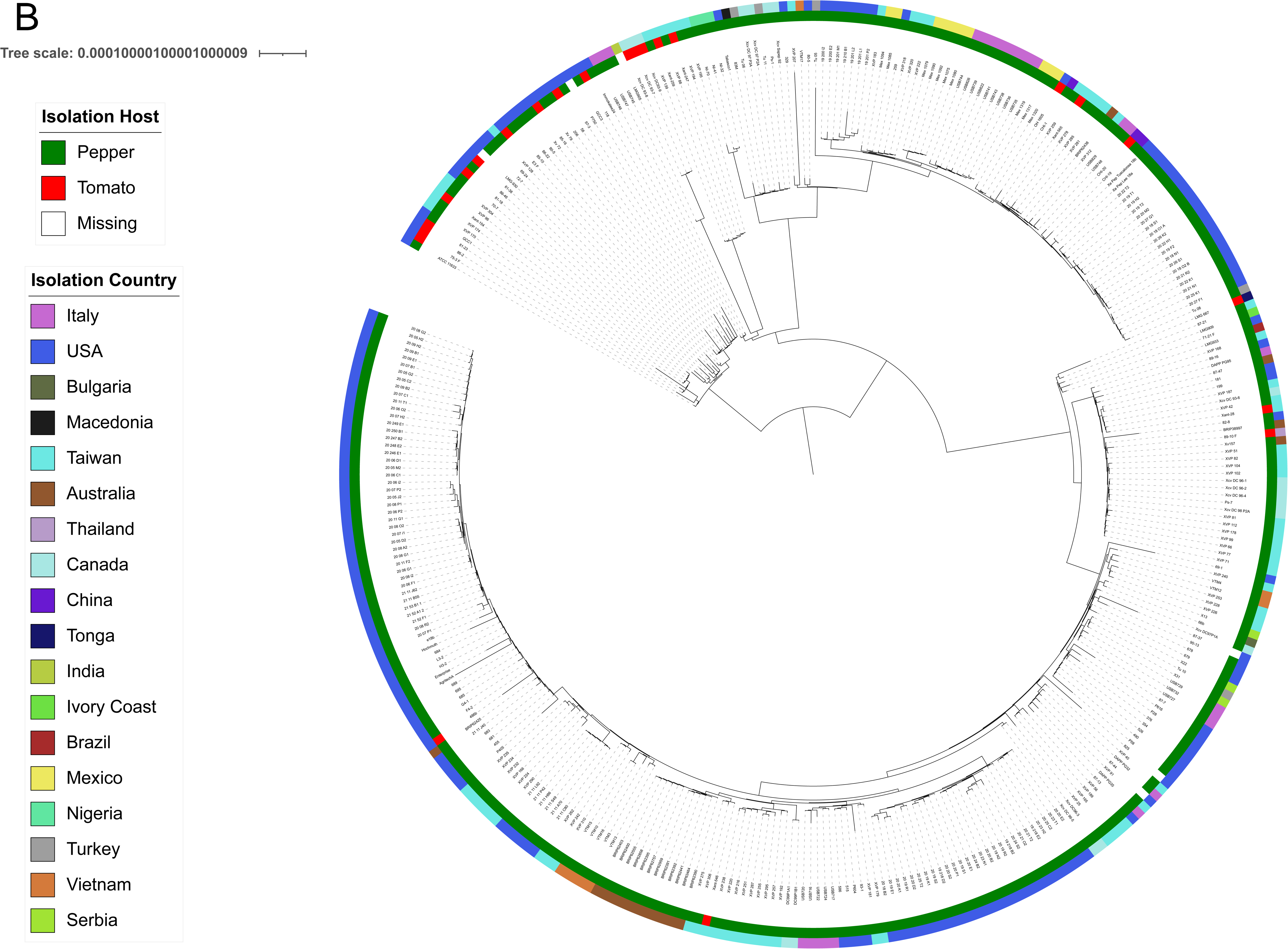
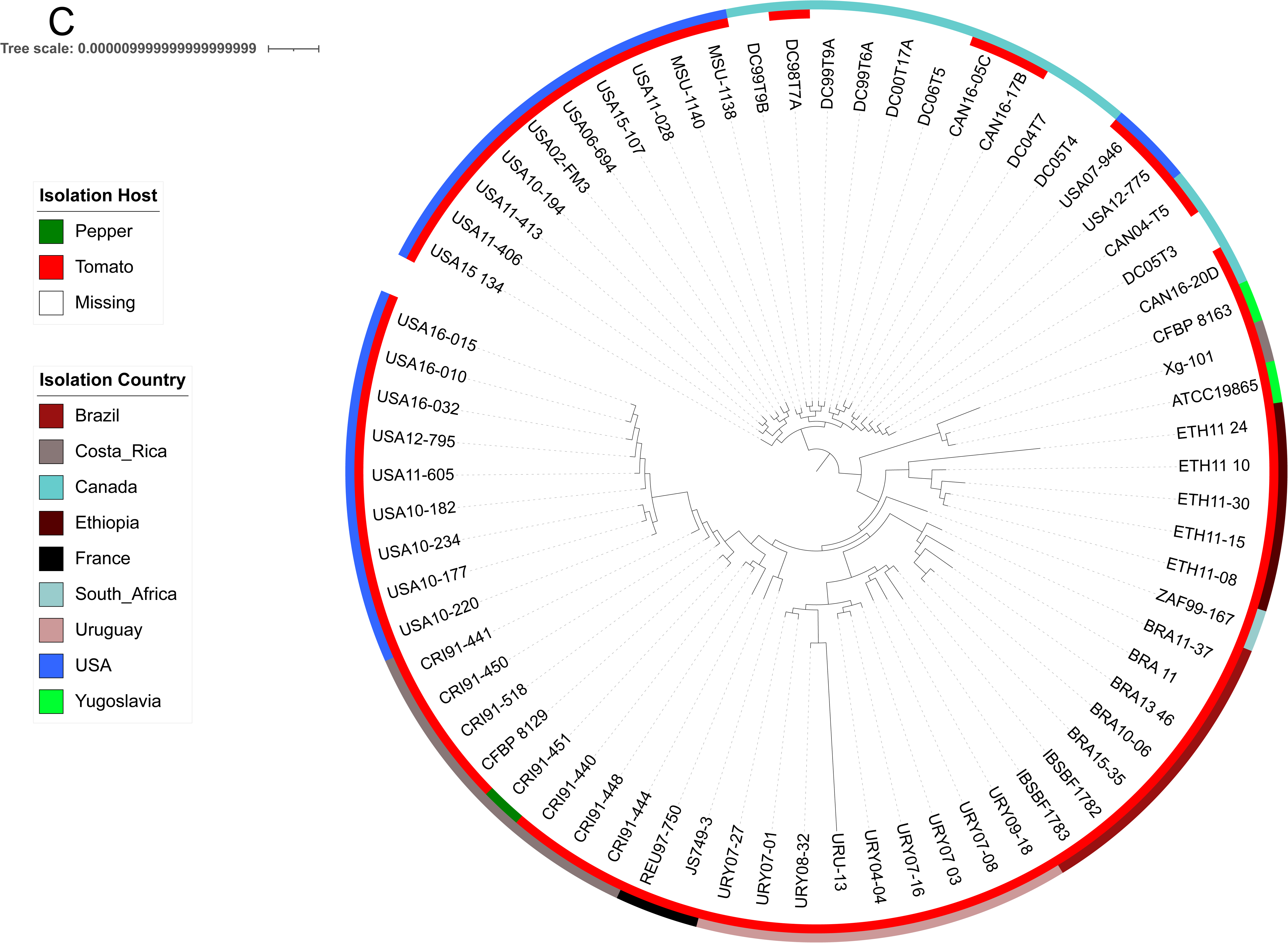
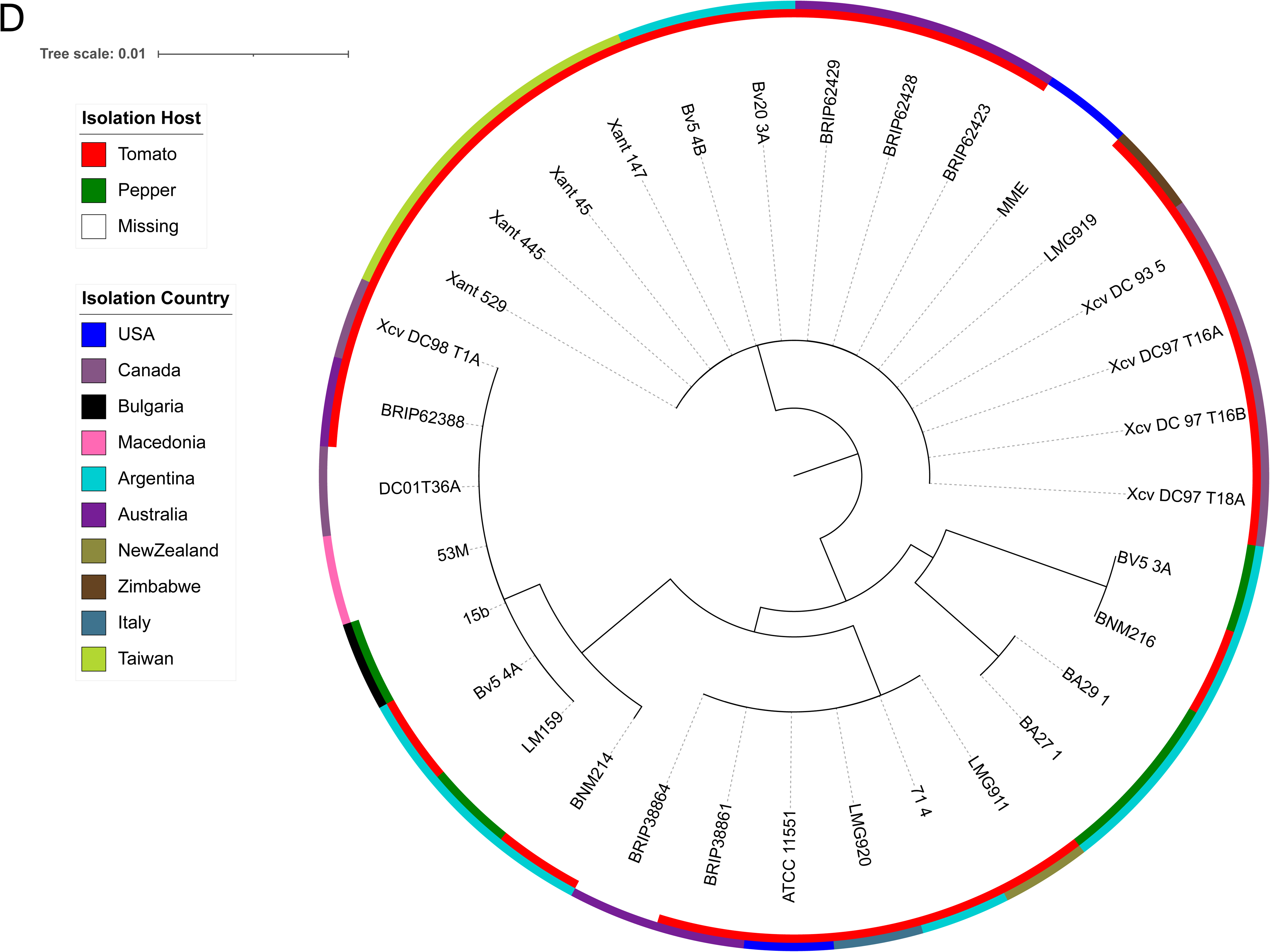
Core-genome maximum-likelihood phylogenetic trees of studied *Xanthomonas* strains. For all panels, the inner ring represents the host, the outer ring represents the country of isolation, and white indicates missing metadata. A. *Xanthomonas euvesicatoria* pv. *perforans* strains from NCBI (n=508) and this study (n=77) based on a core-genome alignment length of 3,949,441 bp. Most strains belong to Lineage A; Lineage B forms a phylogenetically distinct group of 14 strains from Taiwan and Nigeria. B. *X. euvesicatoria* pv. *euvesicatoria* strains from NCBI (n=349) and this study (n=25). Strain LMG918 was omitted to optimize tree visualization. C. *Xanthomonas hortorum* pv. *gardneri* strains from NCBI (n=63) and this study (n=6). D. *Xanthomonas vesicatoria* strains from NCBI (n=33) based on a core genome alignment length of 3,752,908bp.

### Genomic Diversity and Phylogeny of Strains

Phylogenomic reconstruction using 3,806 core genes resolved a deep evolutionary split within the and soft-core effectors as those present in 95% or more but less than 100%. The allelic variation matrices of *Xp, Xe*, *Xg* and *Xv* are depicted in Figure 3 (a-d) and strict-core and soft-core effectors for each pathogen are listed in Table 1. For *Xp*, 42 effectors were identified (Supplementary Table 3a), with 20 strict-core and 8 soft-core effectors (Table 1). Out of 43 total *Xe* effectors (Supplementary Table 3b), *Xe* displayed a repertoire of 11 strict-core and 15 soft-core effectors (Table 1). *Xg* exhibited the highest conservation levels among 27 effectors (Supplementary Table 3c): 21 strict-core effectors and 4 soft-core effectors (Table 1). *Xv* had 17 strict-core and no soft-core effectors (Table 1) out of a total of 26 effectors (Supplementary Table 3d).

**Figure 3.**
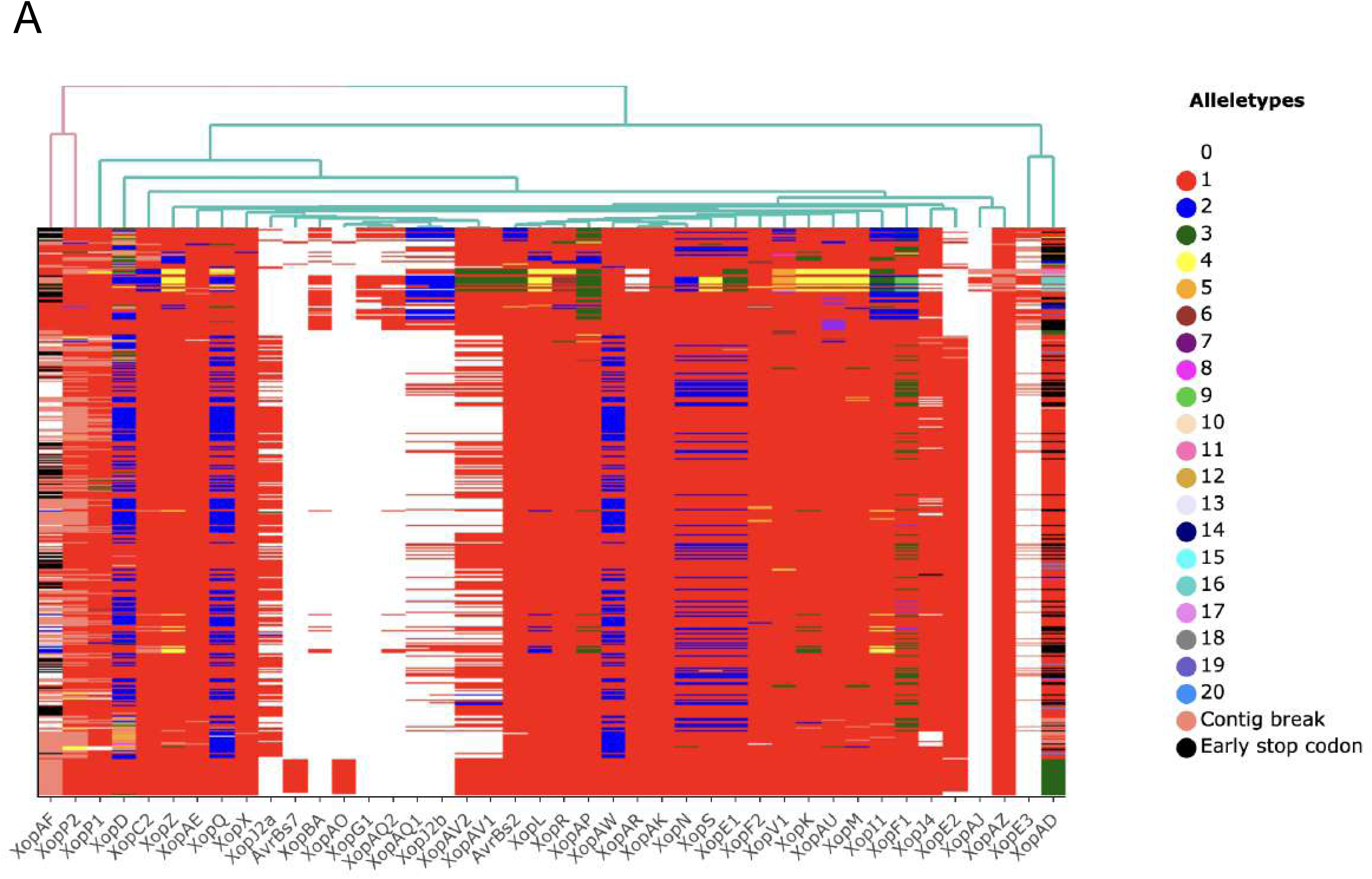
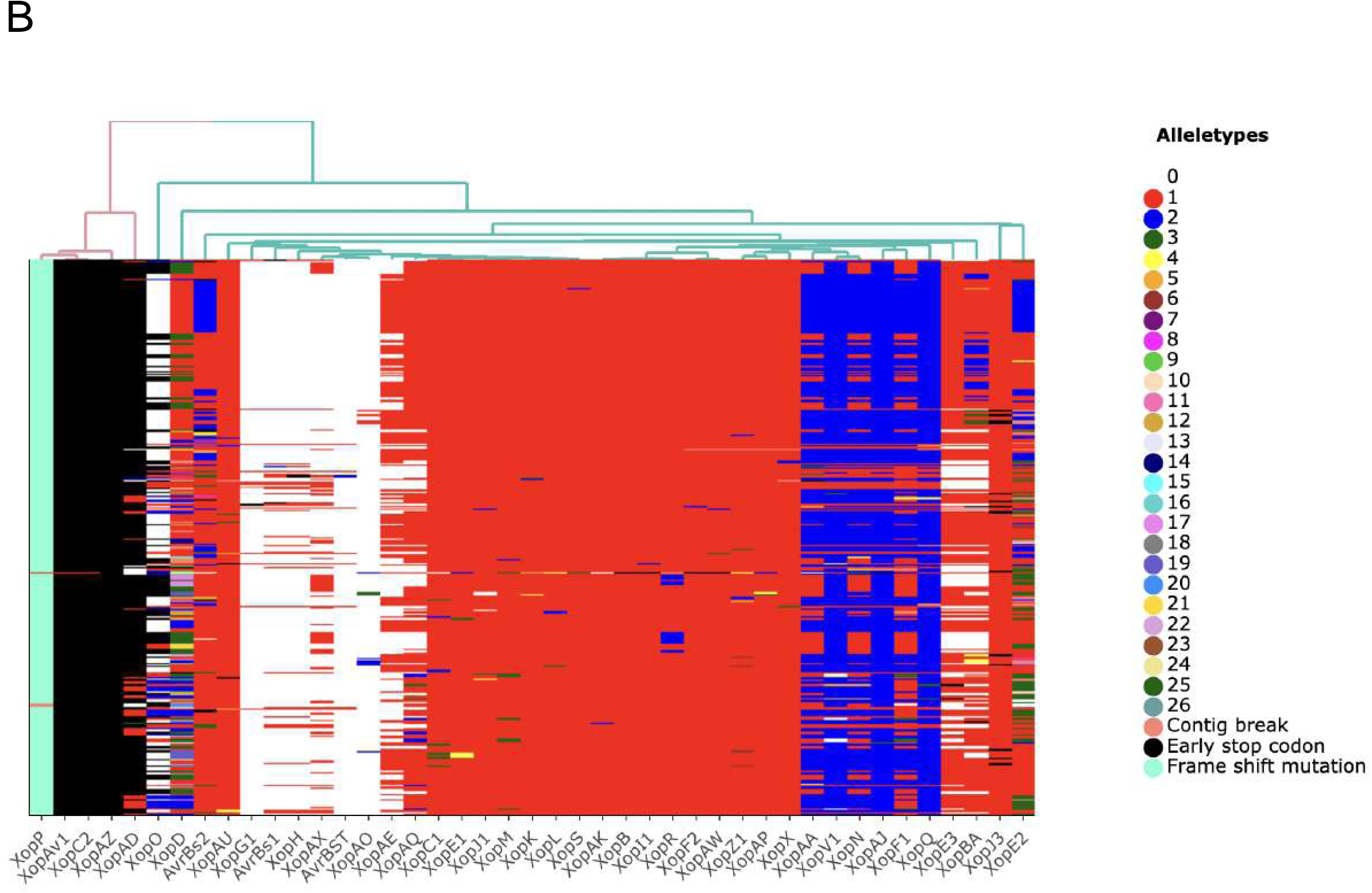
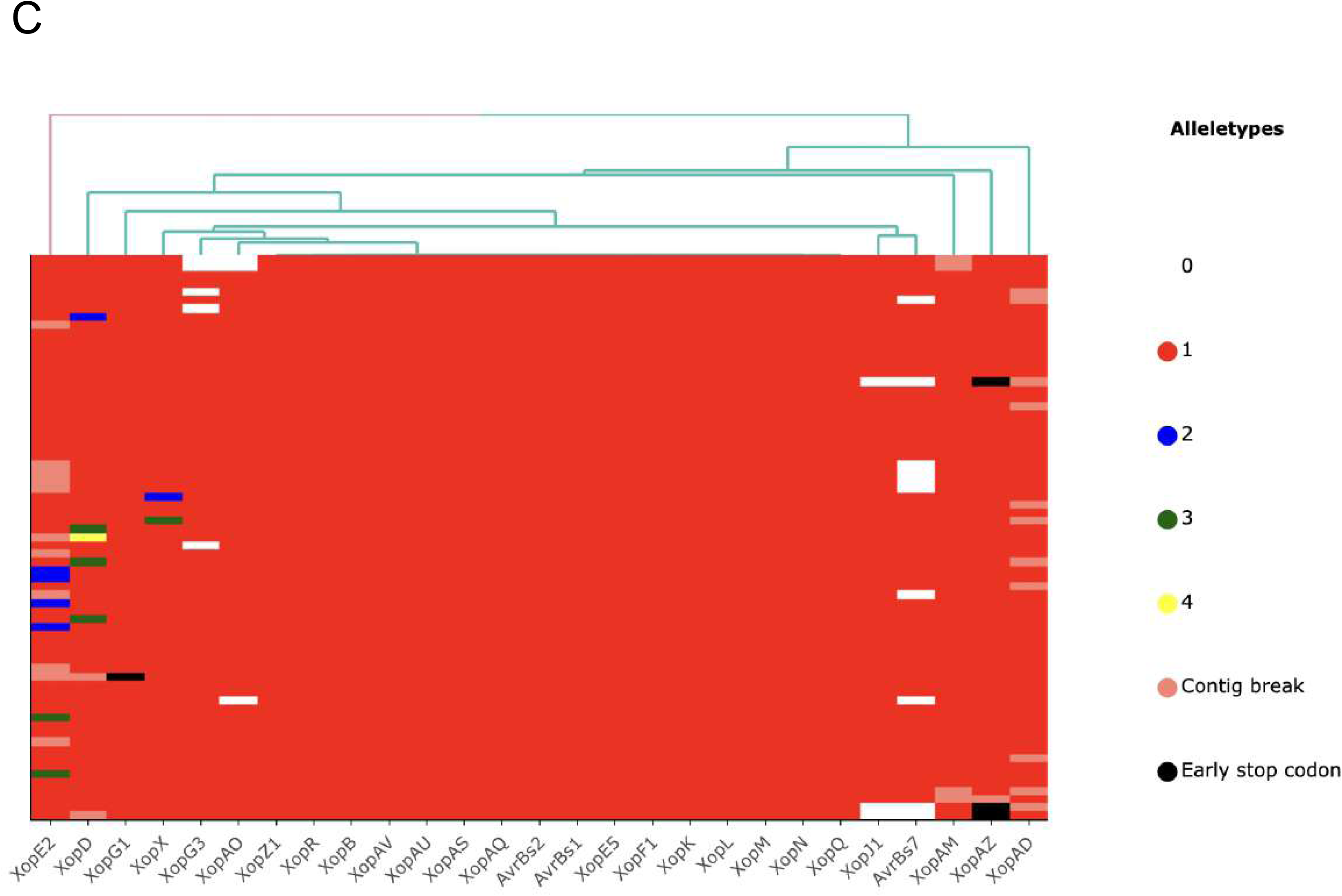
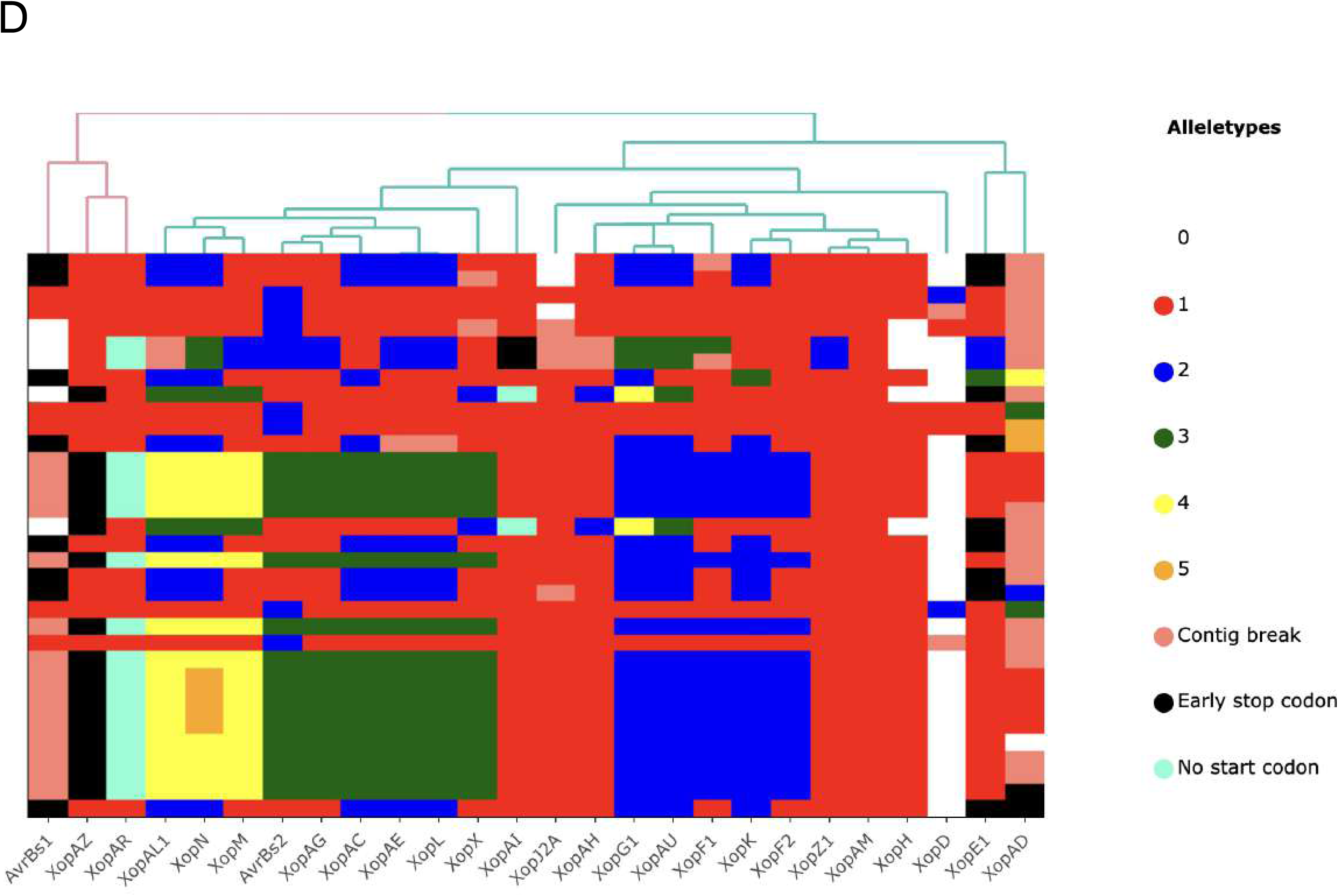
Heat map illustrations of allelic variation in type III effector genes among studied *Xanthomonas* species. For all panels, rows represent individual strains, and columns represent effectors. Numbers and associated colors indicate distinct allele types based on predicted amino acid sequences, with white (0) indicating effector absence. The left-to-right order of effectors was determined via hierarchical clustering of allele types. A. *Xanthomonas euvesicatoria* pv. *perforans*. B. *X. euvesicatoria* pv. *euvesicatoria*. C. *Xanthomonas hortorum* pv. *gardneri*. D. *Xanthomonas vesicatoria*.

**Table 1:**
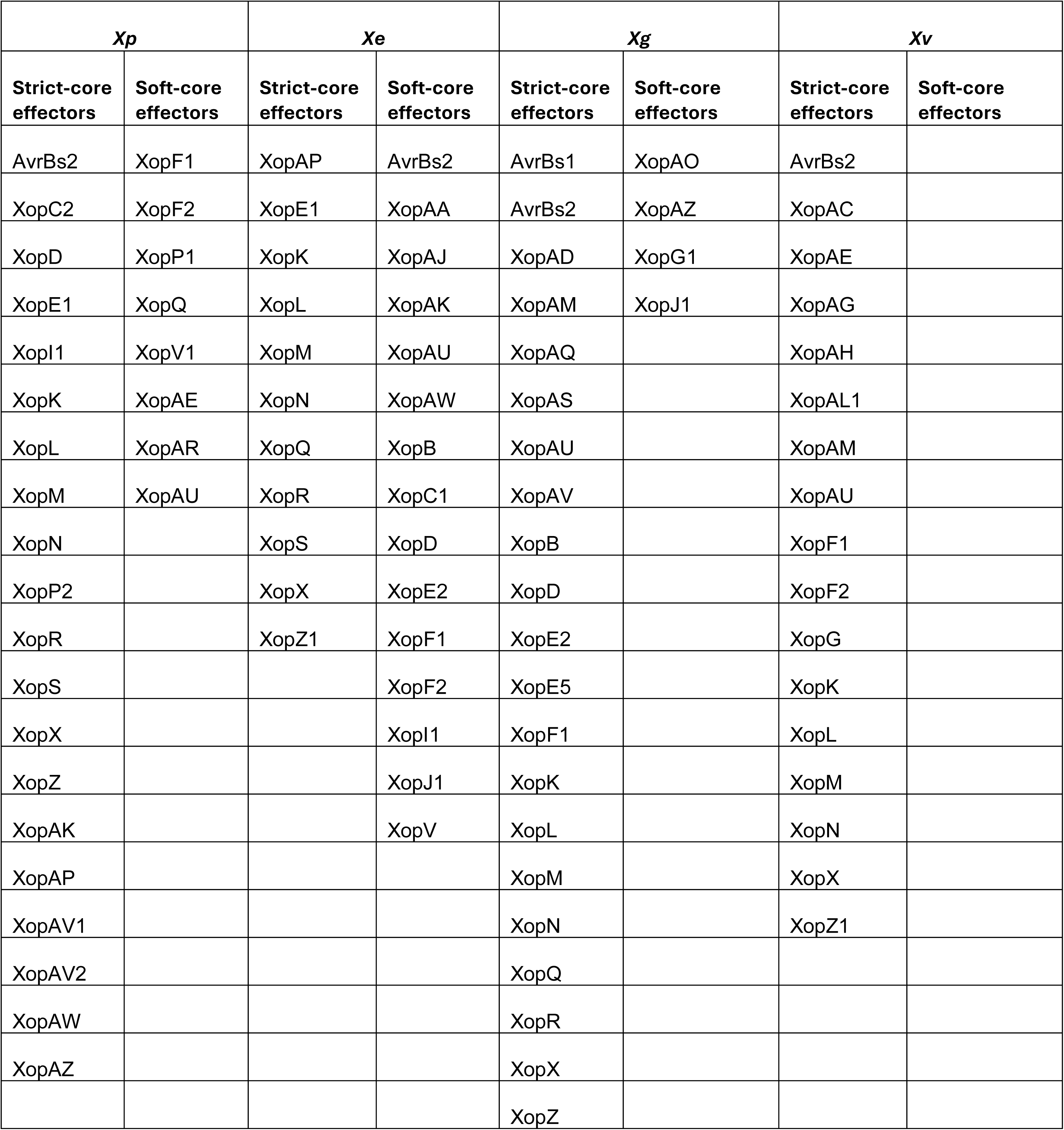
Strict-core effectors were conserved in 100% strains examined and soft-core effectors were conserved in more than 95% but less than 100% strains examined. *Xp, Xe, Xg and Xv* represents *Xanthomonas euvesicatoria* pv. *perforans*, *Xanthomonas* *euvesicatoria* pv. *euvesicatoria*, *Xanthomonas hortorum* pv. *gardneri* and *Xanthomonas vesicatoria* respectively.

| *Xp* |  | *Xe* |  | *Xg* |  | *Xv* |  |
| --- | --- | --- | --- | --- | --- | --- | --- |
| Strict-core effectors | Soft-core effectors | Strict-core effectors | Soft-core effectors | Strict-core effectors | Soft-core effectors | Strict-core effectors | Soft-core effectors |
| AvrBs2 | XopF1 | XopAP | AvrBs2 | AvrBs1 | XopAO | AvrBs2 |  |
| XopC2 | XopF2 | XopE1 | XopAA | AvrBs2 | XopAZ | XopAC |  |
| XopD | XopP1 | XopK | XopAJ | XopAD | XopG1 | XopAE |  |
| XopE1 | XopQ | XopL | XopAK | XopAM | XopJ1 | XopAG |  |
| XopI1 | XopV1 | XopM | XopAU | XopAQ |  | XopAH |  |
| XopK | XopAE | XopN | XopAW | XopAS |  | XopAL1 |  |
| XopL | XopAR | XopQ | XopB | XopAU |  | XopAM |  |
| XopM | XopAU | XopR | XopC1 | XopAV |  | XopAU |  |
| XopN |  | XopS | XopD | XopB |  | XopF1 |  |
| XopP2 |  | XopX | XopE2 | XopD |  | XopF2 |  |
| XopR |  | XopZ1 | XopF1 | XopE2 |  | XopG |  |
| XopS |  |  | XopF2 | XopE5 |  | XopK |  |
| XopX |  |  | XopI1 | XopF1 |  | XopL |  |
| XopZ |  |  | XopJ1 | XopK |  | XopM |  |
| XopAK |  |  | XopV | XopL |  | XopN |  |
| XopAP |  |  |  | XopM |  | XopX |  |
| XopAV1 |  |  |  | XopN |  | XopZ1 |  |
| XopAV2 |  |  |  | XopQ |  |  |  |
| XopAW |  |  |  | XopR |  |  |  |
| XopAZ |  |  |  | XopX |  |  |  |
|  |  |  |  | XopZ |  |  |  |

Allelic diversity analysis highlighted lineage-specific evolution. In *Xp*, XopAD and XopD exhibited the highest diversity with 20 and 11 distinct intact alleles, respectively (Figure 3a, Supplementary Tables 4a and 5a). XopF1, XopV1 and XopP2 had 9, 7 and 7 distinct alleles, respectively. The number of alleles for other *Xp* effectors ranged from 1 to 6. Similarly, *Xe* displayed exceptional polymorphism in XopD, with 26 distinct variants identified (Figure 3b, Supplementary Tables 4b, and 5b). AvrBs2 and XopE2 were other diverse effectors with 9 and 8 alleles, respectively. The number of alleles for other *Xe* effectors ranged from 1 to 6. In contrast, *Xg* effectors were highly stable, with 16 effectors showing identical sequences across all strains (Figure 3c, Supplementary Tables 4c and 5c). XopD was the most variable effector in *Xg* with 4 alleles, whereas other effectors had 3 or less alleles. For *Xv*, XopN and XopAD were the most variable effectors with 5 alleles. The remaining *Xv* effectors had allele numbers ranging from 1 to 4, with only 2 alleles in XopD (Figure 3d, Supplementary Table 4d and 5d).

The divergent *Xp* lineage B (n=14) had unique allelic profiles compared to lineage A. There were alleles specific to the lineage for 9 effectors: AvrBs2, XopC2, XopE1, XopK, XopL, XopM, XopAU, XopAV1 and, XopAV2. There was also variation within the lineage; strains from Taiwan (XVP-273, XVP-272, XVP-271, XVP-270, XVP-267, XVP-265, Xant-477, XTN47 and Xant-592) had specific alleles for XopAD, XopR, XopF1 and XopS and strains from Nigeria (NI1, NI-04, KS3, KS9, and NI-12) had unique alleles for XopAD and XopR (Table 2). Likewise, the divergent lineages in *Xv* contained distinct allelic profiles. One lineage, containing strains collected from 1956 through 2019, demonstrated largely stable allelic profiles despite wide variation in year isolated and country of origin.

**Table 2:**
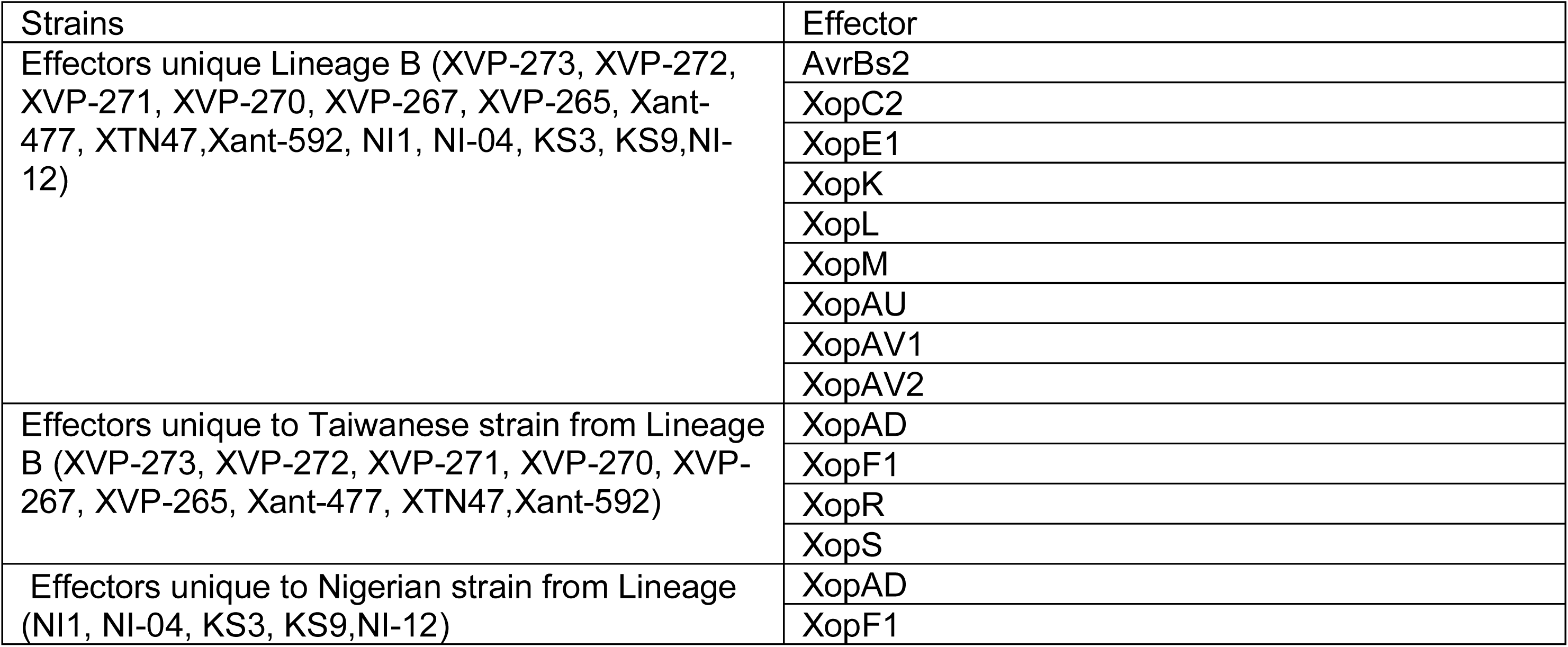
Strict-core effectors were conserved in 100% strains examined and soft-core effectors were conserved in more than 95% but less than 100% strains examined. *Xp, Xe, Xg and Xv* represents *Xanthomonas euvesicatoria* pv. *perforans*, *Xanthomonas* *euvesicatoria* pv. *euvesicatoria*, *Xanthomonas hortorum* pv. *gardneri* and *Xanthomonas ves icatoria* respectively.

### Pseudogenization and Gene Disruption

Disruptive mutations were prevalent in specific effector genes, suggesting ongoing pseudogenization. In *Xp*, early stop codons were abundant in XopAD (n=133) and XopAF (n=98) (Supplementary Table 4a). Within the *Xe* population, XopAZ contained an early stop codon in all examined strains, and XopP1 contained frameshifts in all strains except genetically distinct strain LMG918 (Supplementary Table 4b). Similarly, *Xg* strains exhibited early stop codons in XopAZ and XopG1 (Supplementary Table 4c). In *Xv*, 17 strains had no start codon for XopAR and a partially overlapping 17 strains had an early stop codon in XopAZ (Supplementary Table 4d). Contig breaks were also frequently detected in variable effectors such as XopAD and XopAE across species, which may reflect either structural variation or assembly limitations in effectors with repetitive motifs.

### Effectors conserved across xanthomonads causing BSP/T

To delineate the conserved type III effector (T3E) suites across the four bacterial spot pathogens, we evaluated effector conservation frequencies across our global dataset. Effector conservation profiles were divided into strict-core effectors (present in 100% of strains), soft-core effectors (present in 95% or more but less than 100% of strains), and core effectors (all effectors present in 95% or more strains).

The foundational strict-core T3Es shared across all four pathogens comprised six effectors: XopK, XopL, XopM, XopN, XopX, and XopZ1 (Figure 4A–B). Distinct subsets of effectors were strict-core among combinations of pathogens. Three-way strict conservation was maintained for two additional effectors: XopR was strictly conserved across the *Xe, Xp*, and *Xg* strains, while AvrBs2 was strictly conserved across *Xp*, *Xg,* and *Xv.* Pairwise strict-core comparisons showed *Xg* and *Xv* additionally shared XopAM, XopF1, and XopAU while *Xe* and *Xp* shared an additional three strict-core effectors (XopAP, XopE1, and XopS), while *Xe* and *Xg* shared XopQ as strict-core. No strict-core effectors were shared exclusively between *Xp* and *Xv* nor *Xe* and *Xv*. While *Xp*, *Xg*, and *Xv* each had seven strict-core effectors that were not strict-core in the other pathogens, *Xe* did not have any unique strict-core effectors.

**Figure 4.**
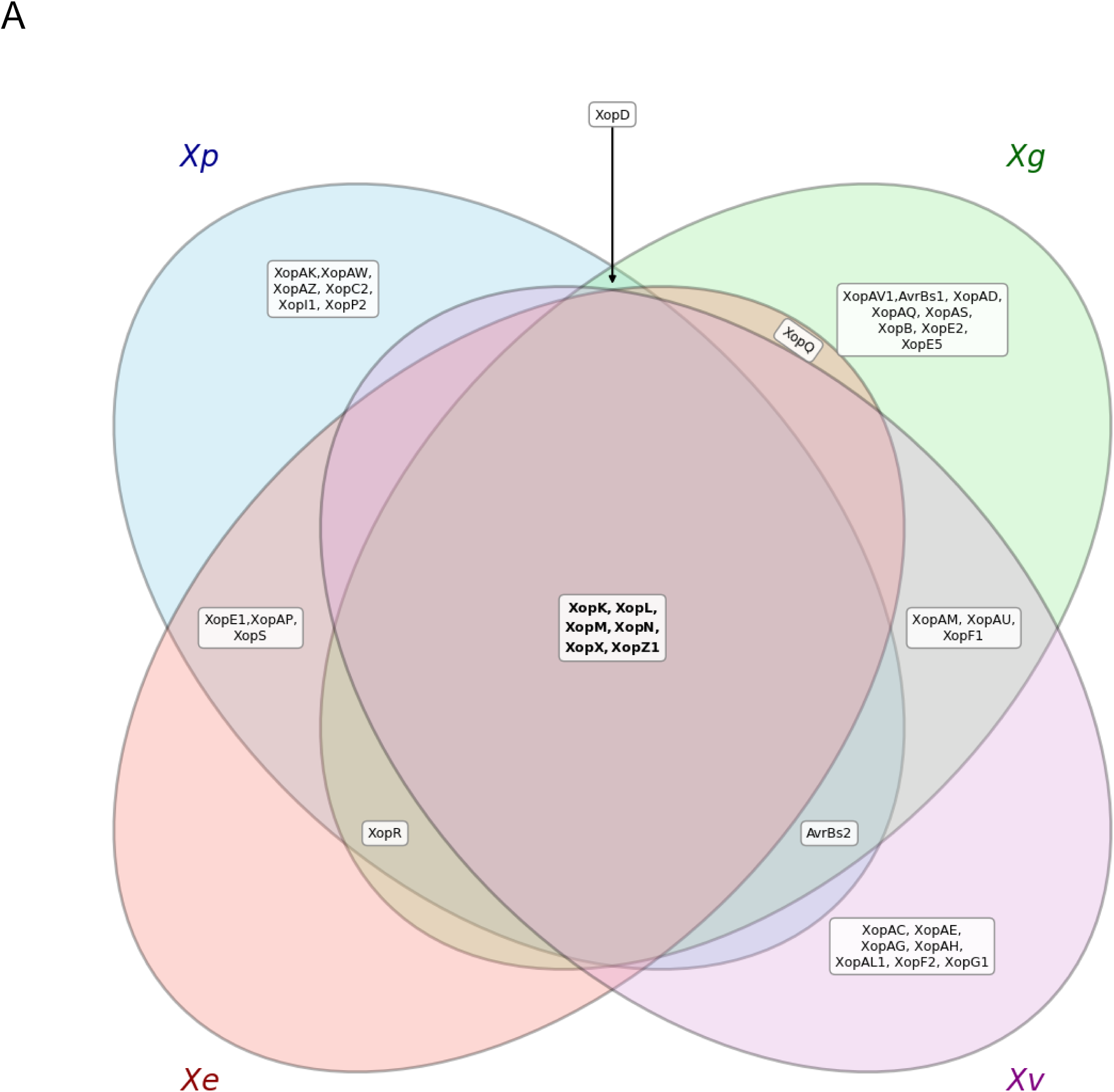
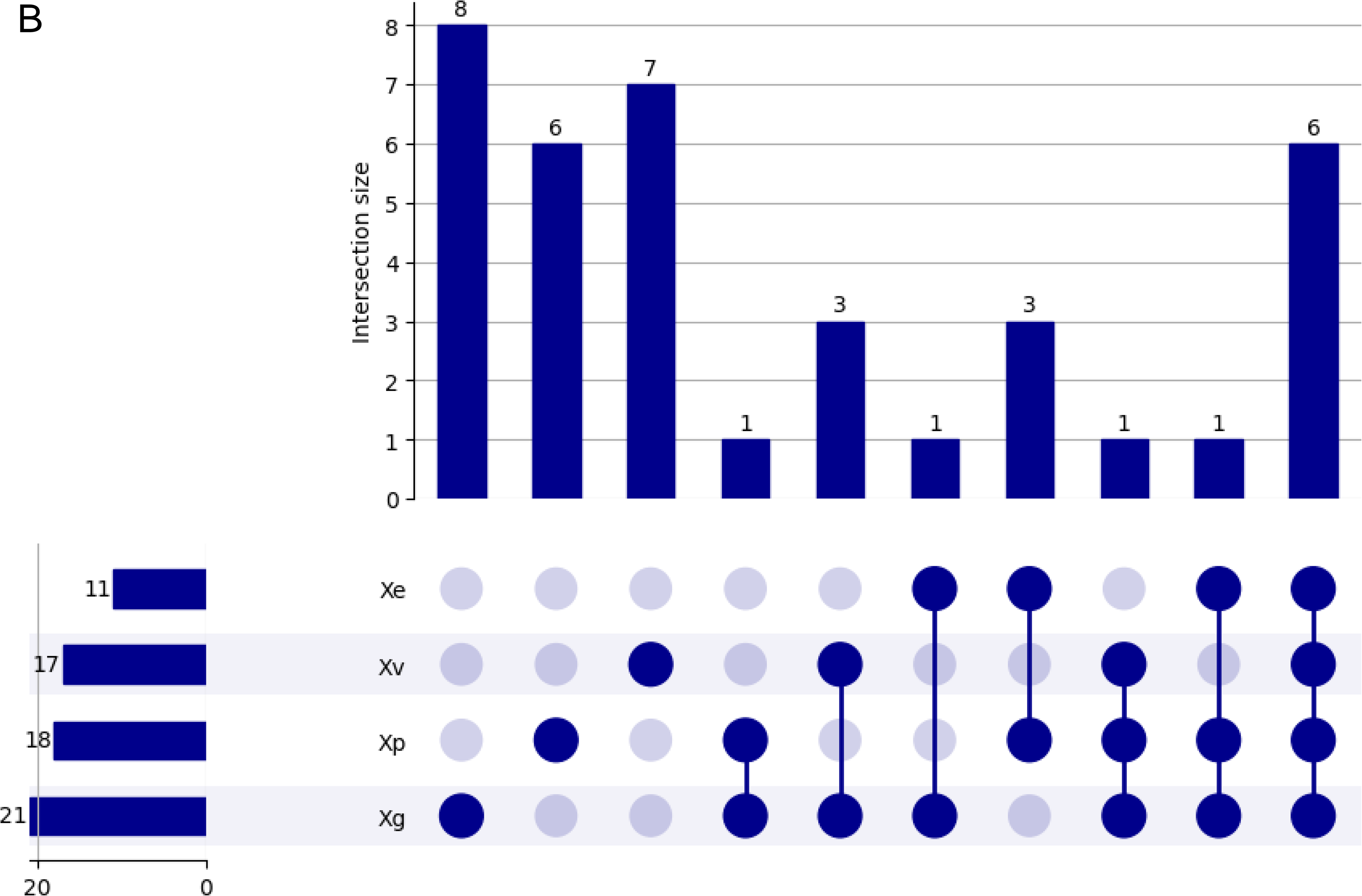
Strict-core (present in100% strains) type III effector (T3E) repertoires in *Xe, Xp, Xg,* and *Xv.* (**A)** Four-way Venn diagram mapping out the shared and unique strict core effector distribution among the four studied lineages. **(B)** Combined UpSet plot and detailed reference matrix indexing the effectors and intersection sizes for every shared or group- specific T3E subset. *Xe, Xp, Xg* and *Xv* represent *Xe*, *Xanthomonas euvesicatoria* pv. *euvesicatoria*; *Xp*, *Xanthomonas euvesicatoria* pv. *perforans*; *Xg*, *Xanthomonas hortorum* pv. *gardneri*; *Xv*, *Xanthomonas vesicatoria*.

Additionally, we looked for soft-core profiles (95% or more but less than 100%) within specific intersection groupings (Figure 5A–B). No soft-core effectors were shared among all four pathogens. Pairwise soft-core intersections revealed that *Xe* and *Xp* shared the largest unique suite of soft-core TSEs, containing XopAU, XopF1, XopF2, and XopV1. Soft-core overlap between *Xe* and *Xg* was limited to XopJ1. No multi-pathogen soft-core components were uniquely captured within any other three-way or pairwise intersections. Pathogen-specific soft-core effectors were heavily skewed toward the *Xe* population, which maintained ten soft-core effectors (AvrBs2, XopAA, XopAJ, XopAK, XopAW, XopB, XopC1, XopD, XopE2, and XopI1). This was followed by *Xp* with four (XopAE, XopAR, XopP1, and XopQ) and *Xg* with three (XopAO, XopAZ, and XopG1). No soft-core effectors were unique to *Xv*.

**Figure 5.**
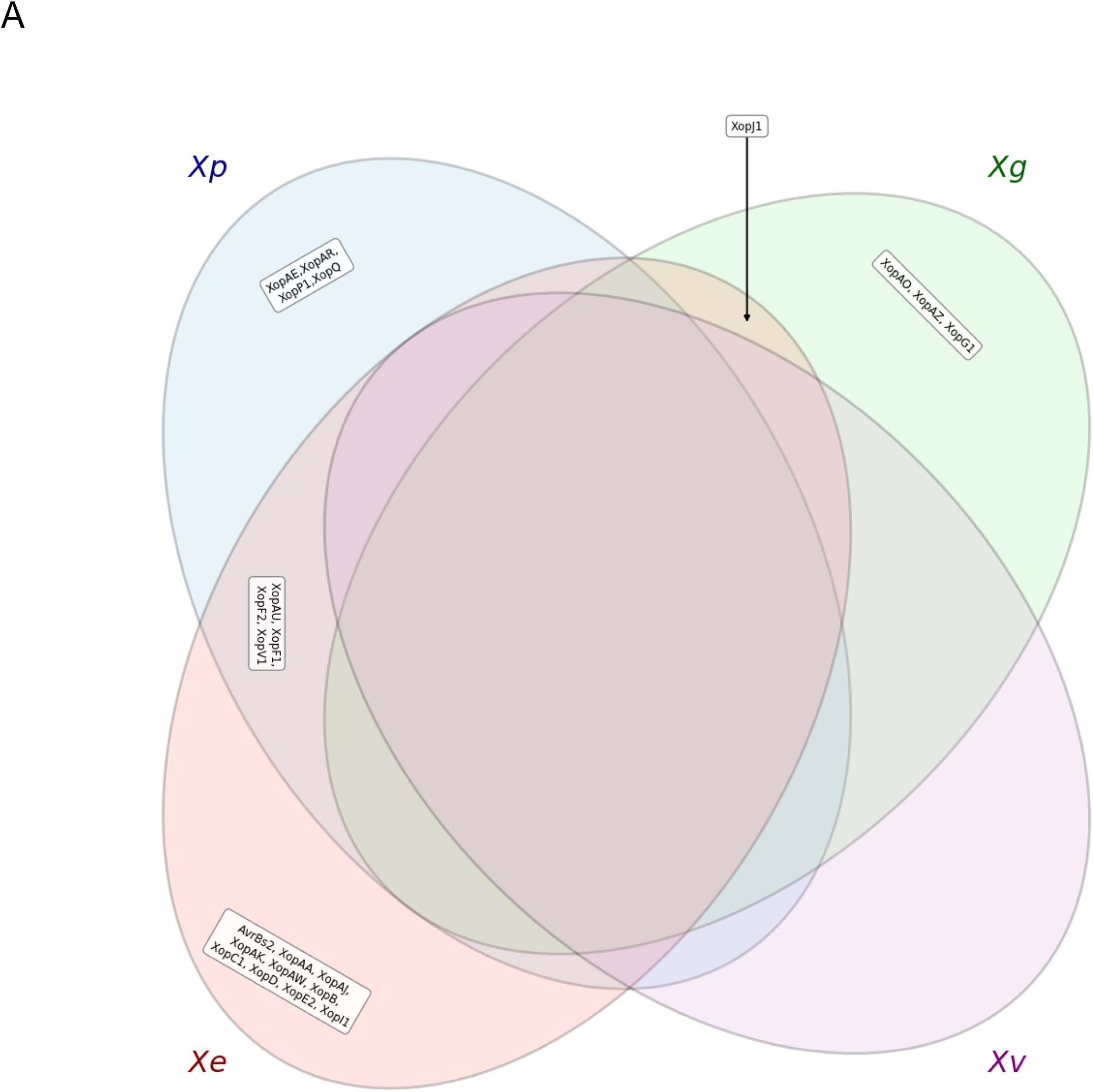
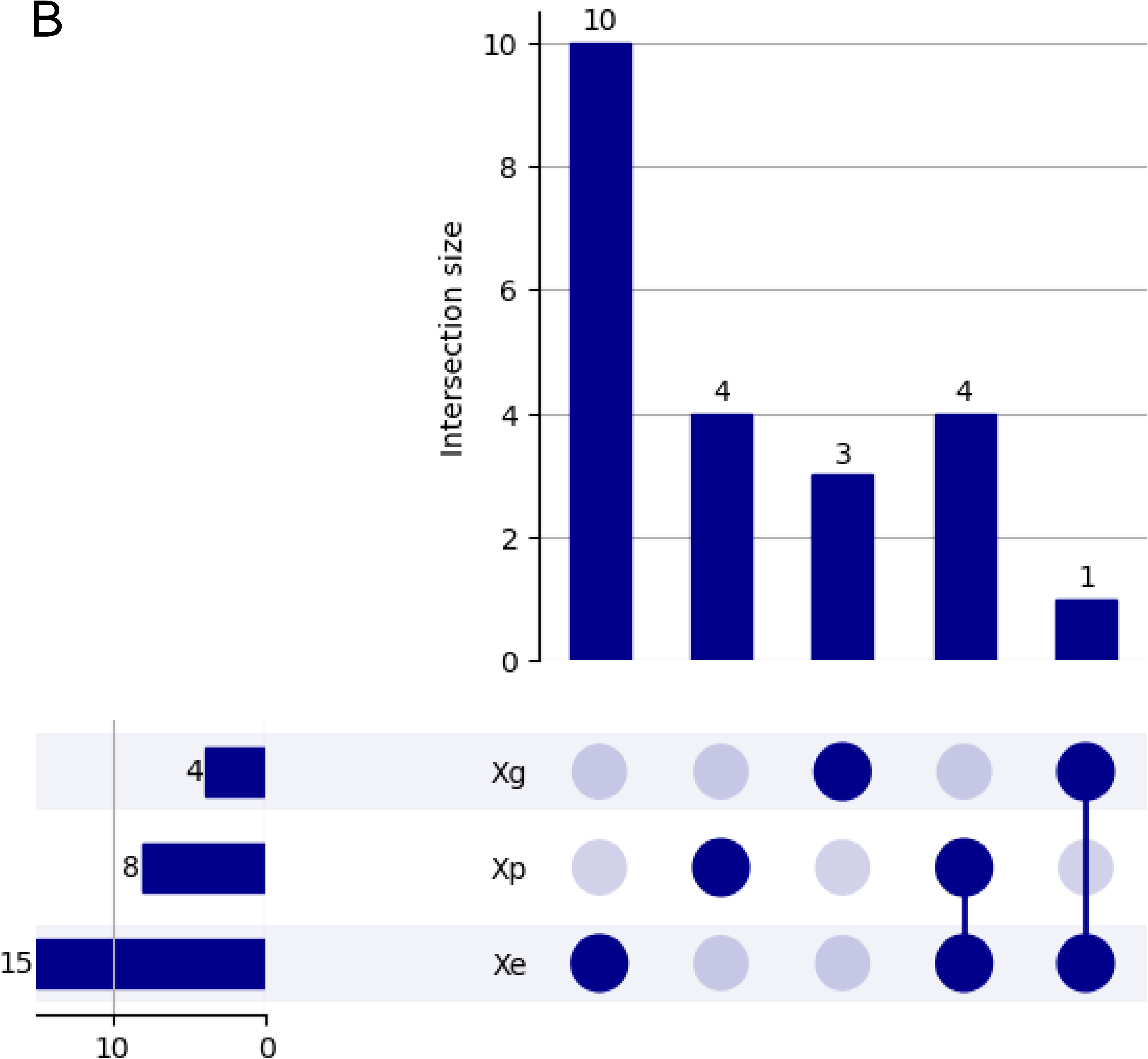
Soft-core (present in ≥95 but <100% strains) type III effector (T3E) repertoires in *Xe, Xp, Xg,* and *Xv.* (**A)** Four-way Venn diagram mapping out the shared and unique effector distributions among the four studied lineages. **(B)** Combined UpSet plot and detailed reference matrix indexing the effectors and intersection sizes for every shared or group- specific T3E subset. *Xe, Xp, Xg* and *Xv* represent *Xe*, *Xanthomonas euvesicatoria* pv. *euvesicatoria*; *Xp*, *Xanthomonas euvesicatoria* pv. *perforans*; *Xg*, *Xanthomonas hortorum* pv. *gardneri*; *Xv*, *Xanthomonas vesicatoria*.

Integrating the strict-core and soft-core datasets revealed the core effector repertoire (95% or more strains) across the bacterial spot pathogens (Figure 6A-B). Under this framework, 9 core effectors were shared across all four pathogens: AvrBs2, XopAU, XopF1, XopK, XopL, XopM, XopN, XopX, and XopZ1. Four three-way shared core effectors were identified for the *Xe*, *Xp*, and *Xg* strains (XopD, XopQ, and XopR) and one for *Xe, Xp*, and *Xv* strains (XopF2). Core effector overlaps were also found in pairwise comparisons for *Xe* and *Xp* (XopAK, XopAP, XopAW, XopE1, XopI1, XopS, and XopV1) and *Xe* and *Xg* (XopB, XopE2, and XopJ1). Smaller subsets of core effectors were shared between *Xp* and *Xg* (XopAZ and XopAV1), *Xg* and *Xv* (XopAM and XopG1), and *Xp* and *Xv* (XopAE). No unique pairwise core effector intersection was observed between *Xe* and *Xv.* Core*-*effectors strictly unique to single pathogens were found across all four clades. *Xg* maintained the highest number of specific core effectors with six (AvrBs1, XopAD, XopAO, XopAQ, XopAS, and XopE5), followed closely by *Xp* with five (XopAR, XopAV2, XopC2, XopP1, and XopP2). *Xv* maintained four unique core effectors (XopAC, XopAG, XopAH, and XopAL1), while *Xe* possessed three (XopAA, XopAJ, and XopC1).

**Figure 6.**
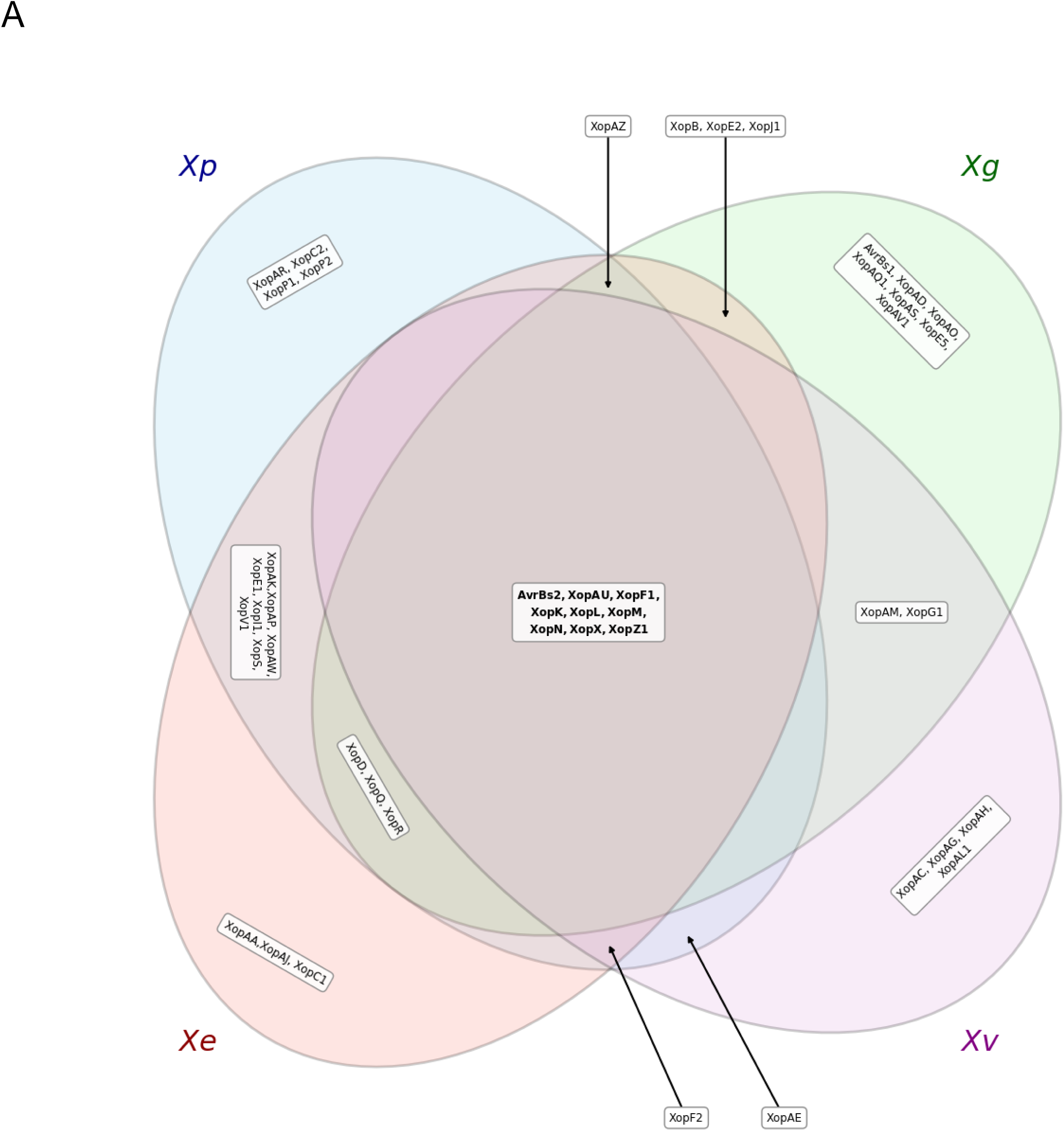
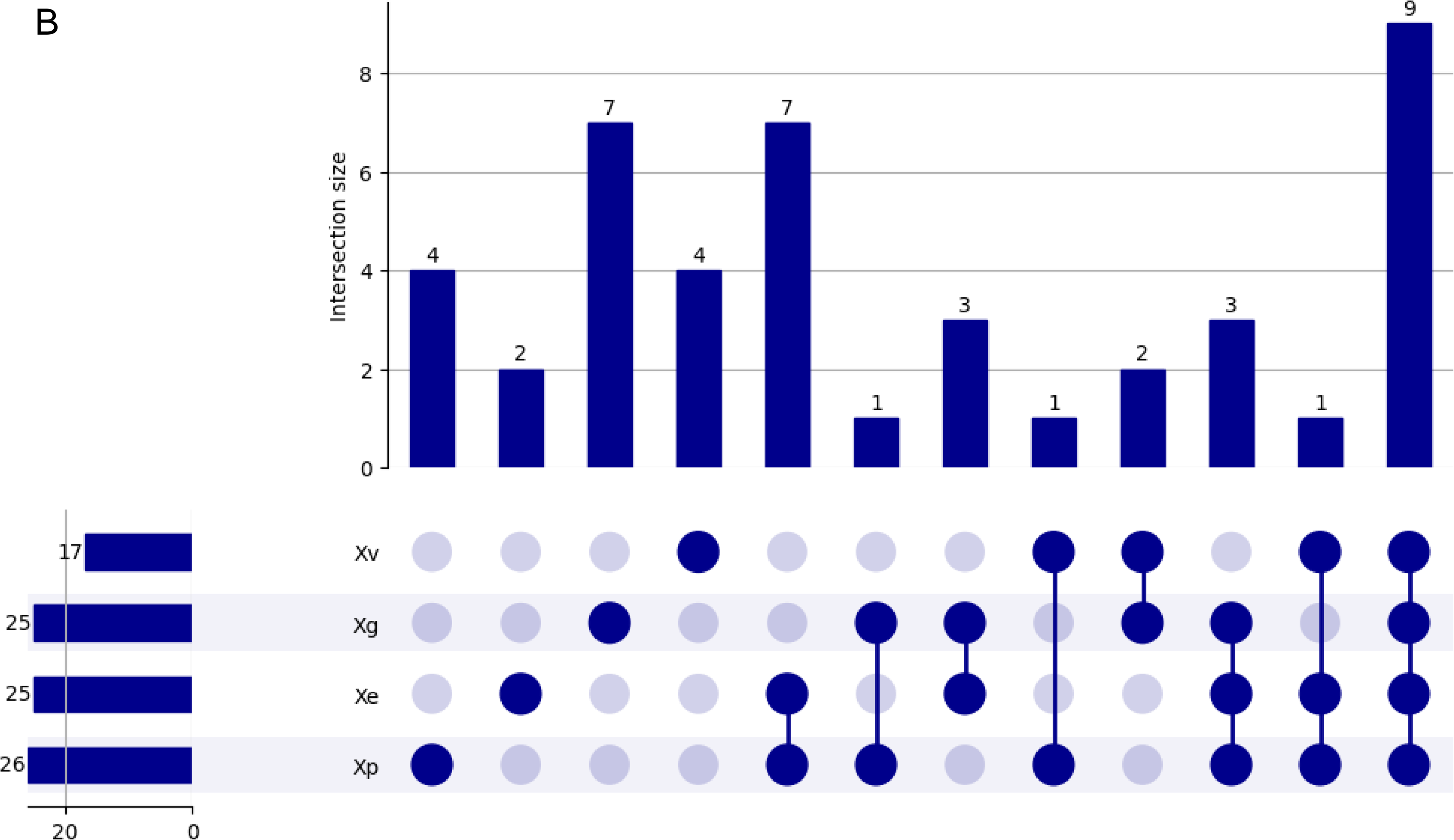
Core (present in≥95 strains) type III effector (T3E) repertoires in *Xe, Xp, Xg,* and *Xv.* (**A)** Four-way Venn diagram mapping out the shared and unique effector distributions among the four studied lineages. **(B)** Combined UpSet plot and detailed reference matrix indexing the effectors and intersection sizes for every shared or group-specific T3E subset. *Xe, Xp, Xg* and *Xv* represent *Xe*, *Xanthomonas euvesicatoria* pv. *euvesicatoria*; *Xp*, *Xanthomonas euvesicatoria* pv. *perforans*; *Xg*, *Xanthomonas hortorum* pv. *gardneri*; *Xv*, *Xanthomonas vesicatoria*.

## Discussion

The comparative analysis of *Xanthomonas* taxa causing bacterial spot revealed six type III effectors that were present in 100% of the 1032 genomes examined: XopK, XopL, XopM, XopN, XopX, and XopZ1. The presence of these six strict-core effectors suggest they perform essential virulence functions required for colonization and pathogenicity in tomato and pepper hosts, likely targeting basal immune responses or fundamental plant physiological processes. Indeed, most of these effectors have been the subject of experimental study. XopN is a virulence factor in *Xe,* because bacterial growth in tomato was impaired when it was deleted in *Xe* strain 85-10 (Kim et al. 2009). XopN was suggested to be an inhibitor of pattern triggered immunity (Popov et al., 2016) by binding to TARK1 (Tomato Atypical Receptor-Like Kinase 1) and TFT1(14-3-3 protein). Following infection, XopN was shown to bind to TARK1 and interfere in TARK1-dependent signaling events, which in turn resulted in the suppression of PAMP-induced gene expression and callose deposition in the host tissue (Kim et al. 2009). Similarly, XopM from *Xe* strain 85-10 functioned as a suppressor of PTI in *Nicotiana* and *Arabidopsis* by mimicking the eukaryotic "two phenylalanines in an acidic tract” (FFAT) motif to bind host Vesicle-Associated Membrane Protein-Associated Proteins (VAPs) at the endoplasmic reticulum and plasma membrane contact sites (Brinkmann et al. 2024). This interaction allowed XopM to dampen early immune responses, specifically reducing the production of Reactive Oxygen Species (ROS), thereby creating a favorable environment for bacterial proliferation. Brinkmann et al. (2024) noted that deletion of XopM in *Xe* did not affect virulence, possibly due to another effector performing redundant functions. XopL from *Xe* strain 85-10 functions as an E3 ligase that directly associates with and disassembles plant microtubules (MTs) (Ortmann et al. 2023). XopL mimics eukaryotic Microtubule-Associated Proteins (MAPs) via a conserved N-terminal Proline-Rich Region (PRR) to facilitate this binding, which is indispensable for induced plant cell death reactions. The direct interaction between the bacterial effector XopL and plant microtubules causes the E3 ligase-dependent depolymerization and collapse of the host cell’s microtubule network. Crucially, this structural positioning on the microtubules is required for XopL to ubiquitinate specific host targets, which ultimately triggers robust plant cell death (necrosis) and suppresses host immunity (Ortmann et al. 2023). XopX, also from *Xe* 85-10, was identified as a virulence factor in tomato and its deletion resulted in a reduction of bacterial growth in tomato leaf apoplast (Stork et al. 2015). Mechanistically, XopX plays]ed a dual role by simultaneously suppressing flagellin-induced ROS generation while paradoxically promoting the accumulation of defense gene transcripts and ethylene production, a process essential for the necrotic symptom development observed in host plants (Stork et al. 2015). XopK was found to be a virulence factor in *Xanthomonas oryzae* pv. *oryzae* (*Xoo*), a major rice pathogen. Deletion of this effector resulted in reduced in planta bacterial growth as well as lesion length in rice. This response was attributed to the E3 ubiquitin-ligase activity of XopK towards OsSERK2 (Qin et al. 2018), a protein responsible for regulation of PTI signaling and resistance to *Xoo* in rice (Hu et al. 2005; Park et al. 2011; Chen et al. 2014). The proven roles of these effectors in the suppression of PTI and/or growth in planta likely explain their strict conservation across the bacterial spot pathogens.

Our population-level analysis also revealed contrasting evolutionary strategies among the four *Xanthomonas* lineages. *Xp* and *Xe* demonstrated expansive total effector repertoires (42 and 43 TSEs, respectively) and core suites comprising approximately two-thirds of the total effectors. Perhaps the most significant finding regarding the *Xp* and *Xe* populations was the extensive allelic variation observed in specific effectors, particularly XopAD and XopD. The taxonomic boundary between *Xe* and *Xp* remains a subject of ongoing debate. For the species delimitation of prokaryotes, average nucleotide identity (ANI) values are frequently used, with values >95% generally representing the same species (Konstantinidis et al., 2006; Goris et al., 2007). Although *Xp* and *Xe* have historically been treated as distinct species (Jones et al., 2004), their >98% ANI has led researchers to classify them as different pathovars of the same species (Constantin et al., 2016). Because of their relatively close phylogenetic relationship compared to the other two bacterial spot pathogens, it is interesting to examine differences in their effector repertoires. While *Xp* and *Xe* shared a large core suite, they also possessed lineage-specific accessory effectors. Specifically, we identified a subset of effectors present in various *Xe* strains but completely absent in the examined *Xp* strains, including AvrBs1, XopAA, XopB, XopC1, XopH, XopJ1, XopJ3, XopO, and XopAX. Conversely, the *Xp* lineage possessed unique accessory effectors—XopJ2b, XopJ4, XopAF, AvrBs7, XopAR, XopP2, XopAQ2, and XopAV2—which are entirely absent from the *Xe* dataset. Effectors such as XopJ1, XopJ2b, XopJ4, and XopJ3 are members of the XopJ family (White et al, 2009) and are known to induce effector-triggered immunity (ETI) in various plant species; specifically, XopJ4 in *Nicotiana* (Schultink et al., 2019; Staskawicz & Schultink, 2024), XopJ3 (AvrRxv) in tomato genotype Hawaii 7998 (Whalen et al. 1993) and XopJ2 in pepper (Sharma et al., 2024). XopAF and AvrBs7 are other proteins responsible for ETI, inducing a hypersensitive response (HR) upon recognition by the resistance loci *Xv3* in tomato (Stall et al., 2009) and *Bs7* in pepper (Potnis et al., 2012), respectively. There were other variants in XopC, XopP, XopAQ, and XopAV that were unique to the *Xp* lineage with closely related homologs in *Xe*. This complementary distribution of homologous effectors strongly suggests functional redundancy. Alternatively, the maintenance of distinct variants from the same effector family in *Xe* and *Xp* may provide specialized fitness advantages tailored to contemporary or past host environments. While the functions of XopAR, XopAA, XopB, XopAX, and XopO remain unknown, their exclusive presence in one taxon or the other suggests they might play specific biological roles that remain to be elucidated. Regardless of individual functionality, the presence of these discrete accessory profiles suggests that, despite their overarching genomic similarity, *Xe* and *Xp* have undergone distinct evolutionary trajectories.

Our analysis of 586 *Xp* strains provides a high-resolution view of effector stability within a single taxon. Despite the phylogenetic divergence into two distinct lineages—a deep-branching group found thus far in Taiwan/Nigeria and a major global lineage—a remarkable degree of conservation was observed in the core pathogenicity arsenal. Twenty effectors were conserved in 100% of the screened strains. This ubiquity indicates that these effectors are indispensable for the lifestyle of *Xp*, regardless of geographic origin or specific host (tomato vs. pepper). However, this stability is contrasted by a subset of accessory effectors that exhibit presence/absence variation. Effectors such as XopF1, XopF2, and XopAE were present in 99% of strains, while XopP1 and XopAR were found in 95%, and XopJ4 in 90%. This variation suggests ongoing adaptation, possibly driven by local environmental pressures or host genotype differences.

*Xg* perhaps represents an alternative evolutionary strategy. While it possesses a small total effector repertoire, almost every effector in the *Xg* arsenal was conserved across >95% of its strains despite temporal and geographic separation in the strains examined resulting in a core effector profile that is comparable in size to the core suites of *Xp* and *Xe*. Like *Xp* and *Xe*, the most diverse effector in *Xg* was XopD (four alleles). Several effectors XopAD, XopZ1, XopAM, XopD, and XopE2 exhibited contig breaks or premature stop codons, suggesting potential pseudogenization or limitations associated with draft genome assemblies. *Xg* was first reported causing bacterial spot disease of tomato in 1950, compared to early 1900s for *Xe* and *Xv*, but its diversification across tomato production regions is minimal compared to *Xp*, which was first reported in 1991.

*Xv*, which was revealed to be a diverse group of strain based on ANI, exhibited a distinct evolutionary trajectory. It possessed a small total effector repertoire, which was similar to *Xg* in total number but with a reduced suite of core effectors. Like *Xp*, distinct core gene lineages contained distinct allelic profiles. The *Xv* strains examined were isolated as long ago as 1956 in Zimbabwe and Italy and as recently as 2019 in Taiwan. While the sample size was small, *Xv* showed stability in alleles across decades and distinct geographic regions within lineages, which suggests a remarkably inflexible genome. Given the dramatic global decline of *Xv* in recent decades, this lack of genomic plasticity within lineages of *Xv* is interesting. The lack of allelic diversity or expansion of its effector arsenal over time might have left it ill-equipped to overcome modern resistant cultivars and agricultural interventions.

The widespread prevalence of disruptive mutations, such as early stop codons and frameshifts across multiple effectors (e.g., XopAD, XopAF, XopAZ, and XopG1), indicates that active pseudogenization is a key feature of *Xanthomonas* evolution. Gene disruption is frequently utilized by bacterial pathogens to quickly shed effectors that trigger host effector-triggered immunity (ETI). The presence of these truncated genes across *Xp*, *Xe*, *Xg,* and *Xv* underscores that gene loss is just as critical to pathogen survival and immune evasion as gene acquisition and allelic variation. This contrasts with the 100% conservation of the core effectors, painting a picture of a pathogen that maintains a rigid, essential virulence backbone while diversifying to persist in the face of host immunity.

## Supporting information

supplemental tabe 1-5

---

**Supplementary Figure 1.**
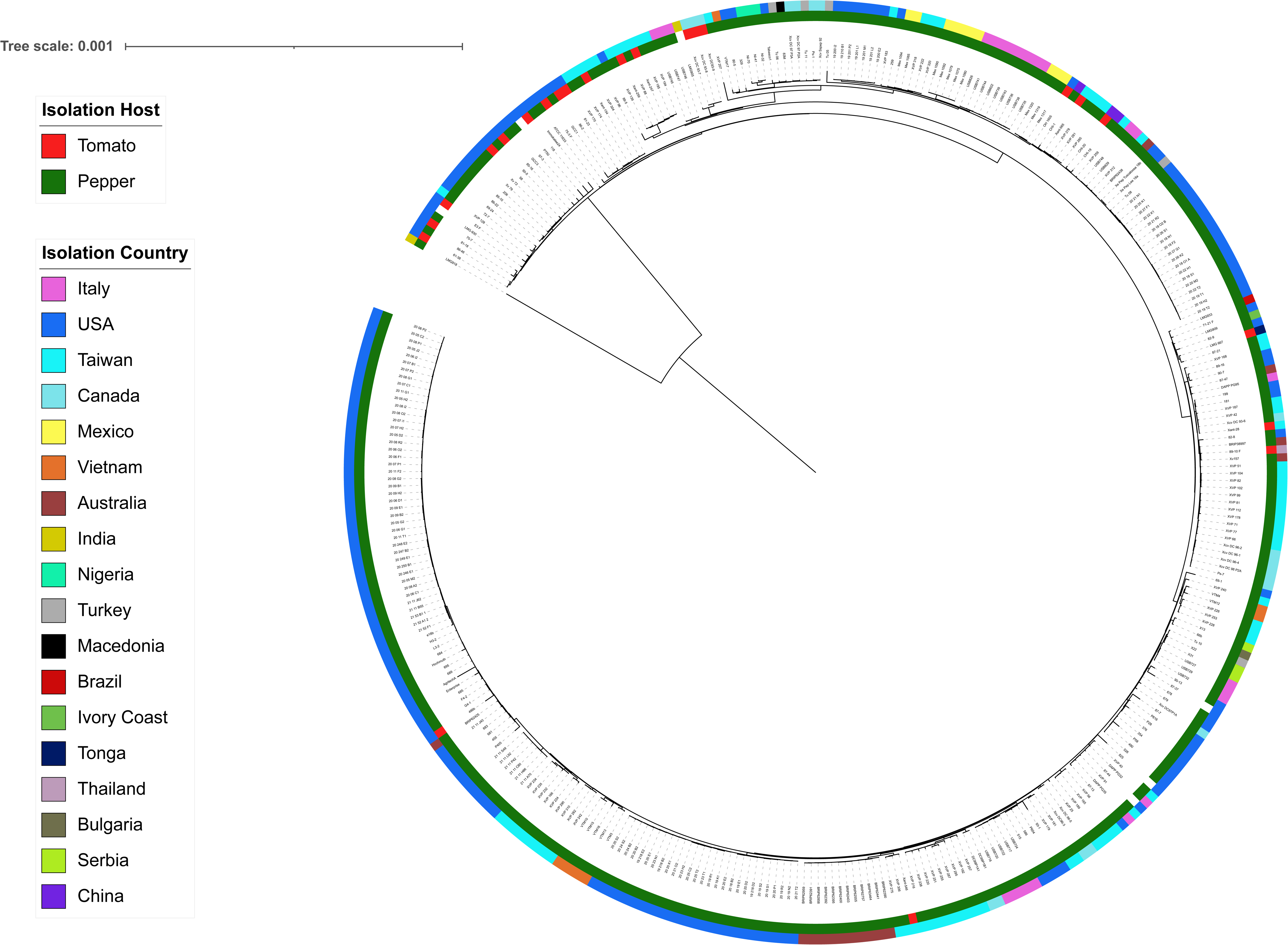
Maximum likelihood phylogenetic tree inferred using core genes of *Xanthomonas euvesicatoria* pv. *euvesicatoria* strains from National Center for Biotechnology Information (n=350) and our collection (n=25). The inner ring represents the host of isolation and outer ring the country of isolation for the respective strains. White indicates missing information. Scale bar is in units of substitutions per site.

